# Late- life shift in caloric intake affects fly longevity and metabolism

**DOI:** 10.1101/2023.05.11.540262

**Authors:** Michael Li, Jacob Macro, Kali Meadows, Dushyant Mishra, Dominique Martin, Sara Olson, Billy Joe Huggins, Brenton Graveley, James Y. H. Li, Blanka Rogina

## Abstract

Caloric restriction (CR) delays the onset of age-related changes and extends lifespan in most species, but how late in life organisms benefit from switching to a low-calorie (L) diet is unexplored. We transferred wild type male flies from a high- (H) to a L-calorie diet (HL) or vice versa (LH) at different times. Late-life HL shift immediately and profoundly reduces fly mortality rate to briefly lower rate than in flies on a constant L diet, and increases lifespan. Conversely, a LH shift increases mortality and hazard rate, which is temporarily higher than in flies aged on a H diet, and leads to shorter lifespan. Transcriptomic changes within 48 hours following diet shift uncover physiological adaptations to available nutrients. Unexpectedly, more abundant transcriptomic changes accompanied LH shift, including ribosome biogenesis, and promotion of growth, which likely contributes to higher mortality rate. Considering that the beneficial effects of CR on physiology and lifespan are conserved across many organisms, our findings suggest that CR interventions in older humans may counteract the detrimental effects of H diets even when initiated later in life.

## Introduction

The incidence of diet-related metabolic disorders such as obesity and type-2 diabetes steadily increases with age. One intervention that may mitigate such disorders is calorie restriction (CR), which is known to improve healthspan and extend lifespan in most model organisms^1–4^. CR leads to physiological adaptations regulated by key nutrient sensing pathways including Insulin/insulin-like growth factor-1 signaling (IIS), sirtuins (SIRT1), and mechanistic target of rapamycin (mTOR) pathways^1,2,5–11^. Among other effects, these pathways regulate body size, growth, reproduction, stress resistance, metabolism, mitochondrial function, autophagy, and lifespan^1–3,5^. Human clinical studies reveal that mild CR or intermittent fasting have many beneficial effects^2,12–16^. However, whether or not an organism can benefit from CR when implemented at an old age is not fully understood, hence more studies into how time of application impact response to CR are needed.

Many studies into CR diets are done in *Drosophila melanogaster* models. Flies aged on a CR diet experience dramatic behavioral, physical, and demographic changes like other organisms^17–20^. Fruit flies have been assayed on a range of different calorie regimens including starvation, intermittent fasting, and overfeeding to determine the effects on stress resistance, spontaneous physical activity, female fecundity, metabolic changes, gut homeostasis, fly health, circadian rhythm and survivorship^17,18,20–27^. Transferring normally fed flies from a standard to a CR diet at young and middle age causes a rapid decrease in mortality rate along with altered gene expression profiles and down-regulation of genes associated with cell growth, and reproduction^19,20,28,29^. Importantly, how late-life shift to CR affect physiology of flies is unclear.

Here we address three key unanswered questions: 1) how late in life shifting flies from a high (H) to a low (L)-calorie diet extends fly life; 2) how shifting from L to H calorie diet late in life impacts survivorship; and 3) what the tissue-specific metabolic and physiological adaptations in response to a shift in dietary caloric content are. We transferred wild type *Canton-S* male flies from H to L diet (HL) or L to H diet (LH) at ages ranging from 10 to 60 days. HL shift reduces age-specific mortality rate to briefly lower rate than in flies on a constant L diet, and increases lifespan even when shifting occurs at a late age. In contrast, LH shift results in an immediate increase in mortality rate that is transiently higher compared to the mortality rate of flies kept on a constant H diet. Head-, thorax-, and abdomen-specific whole genome transcriptional analyses in flies reveal profound modifications following dietary shifts and suggest mechanisms underlying metabolic adaptation associated with lifespan changes. Considering that the beneficial effects of CR on physiology and lifespan are highly conserved across a wide variety of organisms, such examination in fruit flies may be informative for other species.

## Results

### CR has beneficial effects on fly lifespan when applied later in life

To determine whether late-life CR has beneficial effects on fly lifespan, we performed a series of survivorship studies in wild type male *Canton-S* (*CS*) flies. While many studies have investigated the effects of shifting flies from a standard to a CR diet, we shifted flies from a high-calorie (H) diet, 3.0N, to a low-calorie (L) diet, 0.5N. The 3.0N diet has six times higher caloric content compared to 0.5N, which is more clinically relevant to human conditions as metabolic disorders are driven by caloric surplus^18,24^. To determine effects of a late-life shift and to uncover underlying physiological changes, we performed survivorship, metabolic, and transcriptome analyses (Fig. 1A). Flies were aged on a H diet and a subset of flies was transferred to a L diet on day 20 (high-to-low day 20 [HLD20]), 50 (HLD50), and 60 (HLD60). Similarly, a group of flies was aged on a L diet and a subset were shifted to H diet at the same ages (low-to-high day 20 LHD20, LHD50, and LHD60) (Fig. 1A). Males switched from HL at D20 showed a dramatic increase in mean lifespan, and all three age groups have significantly longer maximal lifespan compared to flies kept on a H diet (Fig. 1B). Remarkably, even when flies were switched from H to L at 60 days (HLD60), when only 14 percent of H flies remained alive, maximal lifespan was dramatically extended with HLD60 flies living 35.2% longer than flies on a H diet (Fig. 1B, Table S1,2). Surprisingly, comparing mortality kinetics uncovers that HL and LH shifts lead to immediate and long-term effects. HL shift at ages 20, 50, or 60-days, resulted in an acute and short-term decrease in age-specific mortality rate, which was briefly lower than of flies on a constant L diet but soon became similar to flies on constant L diet (Fig. 1D-F). Conversely, there was an immediate increase in short-term risk in LH-shifted flies at all three ages (LHD20, LHD50, LHD60), which was briefly higher and then became similar to that of flies on a constant H diet (Fig. 1G-I). However, the LHD50 shift decreases maximal survivorship below that observed in flies on a constant H diet uncovering that shifting LH later in life is more detrimental than living on H diet the whole life, while HL shift is profoundly beneficial even at old age (Fig.1I).

**Figure 1:**
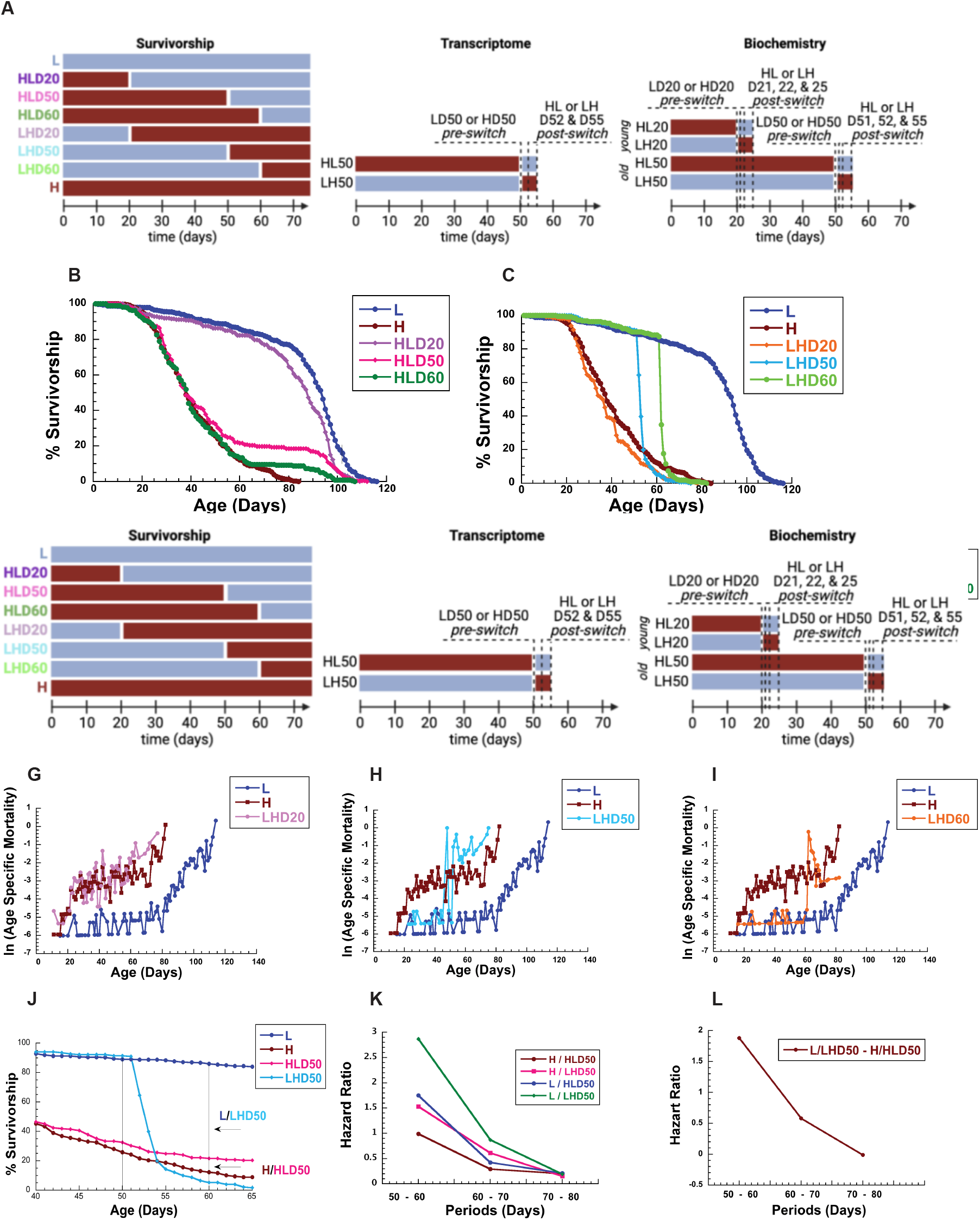
Shifting diets has profound effects on male lifespan and mortality: **A)** Experimental diagram: male flies were kept on a low calorie (L), or a high calorie diet (H) and shifted from H to L diet at day 20 (HLD20), 50 (HLD50), or 60 (HLD60), or from L to H diet on day 20 (LHD20), 50 (LHD50), or 60 (LHD60). Transcriptomes were determined in flies kept on a L or a H diet at day 50 (D50) and 2 and 5 days after H to L (HLD52, HLD55) or L to H (LHD52, LHD55) shift. Metabolism was determined in homogenate of flies kept on a low calorie (L) or a high calorie diet (H) at 20 (LD20, HD20) or 50 (LD50, HD50) days, and in flies 1, 2, or 5 days after shifting from H to L diet at day 20 (HLD21, HLD22, HLD55), or day 50 (HLD51, HLD52, HLD55), or from L to H diet on day 20 (LHD21, LHD22, LHD25), and day 50 (HLD51, HLD52, HLD55). **B-E)** Survivorships (B,C) and mortality rates (D-I) of male flies shifted from H to L at day 20 (HLD20), day 50 (HDL50), or day 60 (HLD60) (B,D-F) or from L to H diet on day 20 (LHD20), day 50 (LHD50), or day 60 (LHD60) (C,G-I). Survivorship curves and mortality rate were analyzed by long-rank test JMP16 program. **J-L)** HL shift decreases, and LH shift immediately increases hazard ratio. HR = 1: no effect, HR > 1: increase in hazard, HR < 1: decrease in hazard (see methods for details). **J)** Survivorship of male flies between 40–65 days to illustrate dramatic changes in survivorship between flies on a constant L or H calorie diet, and flies shifted from a H to L (HL) or L to H (LH) diet at D50, **G)** Hazard ratio and **H)** hazard ratio difference between L and LHD50 or H and HLD50 ratios.

### Diet shift has profound effects on hazard ratio

Cox regression analysis was used to calculate differences in hazard ratios (HR) between flies on a constant L or H diet (control) and LH- or HL-shifted flies (experimental), (Fig. 1J-M)^30^. The data were binned in 10-day intervals to evaluate how diet shifting later in life affects lifespan. HLD50 shift decreases HR illustrated by a significant decrease in H/HLD50 ratio of 0.29 (p=0.02) and 0.2 (p=0.008) in male flies 20 and 30 days after shifting, respectively (Fig. 1K, Table S3). Remarkably, the HLD50 shift decreases HR when compared to HR of flies on constant L diet (Fig. 1K) illustrated by HR L/HLD50 of 0.21 (p=0.01) and 0.25 (p=0.02) 30 and 40 days after shifting, respectively (Fig. 1K, Table S3). In contrast, high transient increased mortality rate and short-term risk of death were observed in LHD50 and LHD60 flies immediately following shifting compared to flies on a constant H diet. This is illustrated by the death of 186 LHD50 (80%) flies during the first 10 days after shifting compared to 22 (5.7%) on a constant H diet (Fig. 1J). This is reflected in 1.53 H/LHD50 HR (p= 0.005) and increased HR of 2.87 L/LHD50 (p<0.001) when compared to L flies during the first 10 days after shift (Table S3A). Shifting from L to H at D60 resulted in a similar trend in HR, with the highest increase in HR L/LH followed by H/LHD60, L/HLD60, and the smallest H/HLD60 (Table S3B). While LH-shifted flies at D20 have only a small increase in HR (1.19), their lifespan is shorter compared to flies on a constant H diet (Table S3C). Next, to examine physiological changes associated with shifting diets, we performed metabolic and transcriptome analyses.

The beneficial effects of late-life shift were confirmed in a second set of shifting experiments. *CS* male flies were shifted HL or LH at day 10, 20, 30, 40, and 50. Both shifts affect mean and maximum life spans and mortality rates at all ages (Fig. S1, Tables S4,5). Male flies maintained on constant H diet had 18.4 and 12.6 percent shorter mean and maximal lifespans, respectively, compared to HLD50 flies. Conversely, LH shift (LHD50) resulted in an immediate increase in mortality rate with 28.6 and 17.6 percent shorter mean and maximal lifespan, respectively, compared to flies on a constant L diet (Table S5).

### Diet shift induces striking tissue-specific changes in transcription

To investigate mechanisms underlying profound changes in survivorship during late life diet shifts, we determined tissue-specific transcriptional changes in the flies. RNA was isolated from the heads, thoraces, and abdomens of HD50 or LD50 male flies, and in flies 2 and 5 days after diet shift (Fig. 1A). We determined RNA-Seq profiles using three biological replicates at each time point. A sample-to-sample distance heatmap shows tissue type to be the driving force for distance, indicating distinct transcriptional profiles across tissue types (Fig. S2A). This is corroborated by transcriptome-wide principal component (PC) analysis, which indicate that biological samples from the same tissues cluster together and are clearly distinct (Fig. 2A). PC analyses of samples from the head or abdomen indicate that samples segregate along PC1 and PC2 primarily by diet. Head and abdomen samples isolated 2 or 5 days after shifting to a L diet, cluster with samples on constant L diet (Fig. 2A). Samples from the thorax show greater variance along PC1 but smaller PC2 variance (Fig. 2A). A heatmap of unsupervised clustering of differentially expressed (DE) genes from all samples and heatmaps of samples separated by LH and HL shifts, confirm clustering at the diet and tissue level, respectively (Fig. 2B, Fig. S2B). Gene expression in biological samples from the head, thorax, or abdomen cluster together based on diet before shifting (LD50 or HD50), while gene expression collected on days 52 and 55 cluster together and with samples from the diet the flies are shifted to (Fig. 2C). Taken together, two days after shift transcriptional responses are closer to transcriptomes of samples from the new diet than to original diet.

**Figure 2:**
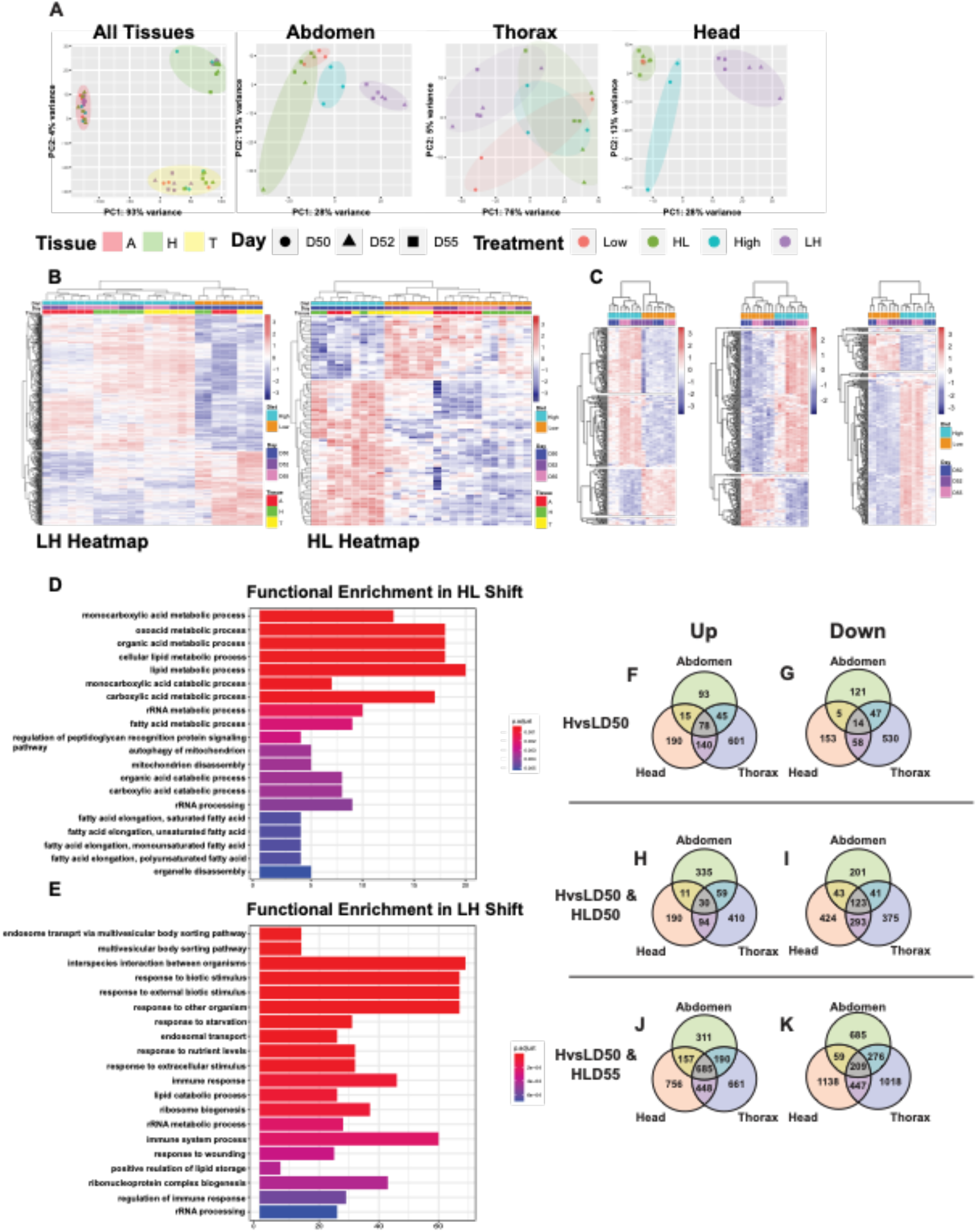
Shifting diet induces striking tissue-specific changes in transcriptome: **A)** PCA plots of all tissues or specific tissues, marked Abdomen, Thorax, or Head, of differentially expressed (DE) genes at high (H) or low (L) diets at day 50 (D50) before, and 2 and 5 days after H to L (HLD52, HLD55) or L to H (LHD52, LHD55) shifting. **B**,**C)** Heatmaps of all (B) or tissue-specific (C) DE genes between flies on H or L diets at day 50 (D50) before and 2 and 5 days after HL (HLD52, HLD55) or LH (LHD52, LHD55) shift. **D**,**E)** Functional enrichment analysis of DE genes in heads, thoraces and abdomens associated with HL (D) and LH (E) shift. **F-K)** Venn diagrams show upregulated (Up) (F,H,J) or downegulated (Down) (G,I,K) DE genes between L and H diet from head, thorax, and abdomen at day 50 before shifting (HvsLD50) (F,G) or 5 days after HL shift (HvsLD50 & HLD55) (H,I) or LH shift (HvsLD50 & LHD55((J,K). FDR threshold: 0.05, logFC threshold: 0.

### Functional enrichments in HL shift reveal strong fatty acid metabolism

We used DESeq2 to quantify differential expressed (DE) gene in head, thorax, and abdomen samples obtained from flies before and after shifting^31,32^. To identify specific pathways associated with HL and LH shift across all tissues we performed functional enrichment analysis of each shift separately (Fig. 2D,E). This analysis revealed specific KEGG metabolic processes that mediate HL adaptation across all tissues including monocarboxylic acid metabolic and catabolic processes, carboxylic metabolic processes, cellular lipid metabolic processes, and rRNA processing. Strong functional enrichment was associated with FA metabolism, including FA elongation of saturated, unsaturated, monosaturated, and polyunsaturated FAs.

These results suggest a shift in metabolism, with flies on a low diet utilizing FAs as preferred means for substrate oxidation, which is consistent with data found in flies, rodents, and humans^2–5,20,28,33–37^.

### Functional enrichments in LH shift reveal increased ribosomal biogenesis and lipid storage processes

Increase in nutrients provided by a H diet leads to positive regulation of proliferation, and lipid storage (Fig. 2E)^38^. Such processes were identified in LH shifted flies by functional enrichment analysis and include ribosome biogenesis, rRNA processing, ribonucleoprotein complex biogenesis, rRNA metabolic processes, and positive regulation of lipid storage, all processes that demand high energy investment. Further, regulation of the immune response was another significantly enriched pathway, which is reported to be activated under high calorie diet conditions (Fig. 2E)^39^.

### Dynamics of DE genes associated with H and L diets and shifts

To search for common and tissue-specific pathways associated with CR, we compared head-, thorax-, and abdomen-specific DE genes between H and L (HvsL) diets before shifting (HvsLD50) (Fig. 2F,G). A Venn diagram shows only 78 commonly upregulated and 14 commonly downregulated DE genes across the three tissues (Fig. 2F,G). The largest number of unique genes are found in the thorax (up=601, down=530), while a similar number of unique genes are found in the head (up=190; down=153) and abdomen (up=93; down=121) (Fig. 2F,G,). Surprisingly, comparison of head, thorax, and abdomen DE genes before- and 5 days post-shift (HvsLD50 & HLD55) reveals only 30 commonly upregulated and 123 downregulated genes across the three tissues (Fig. 2H,I). Most of these genes are involved in metabolic processes.

In contrast, LH shift increases the number of commonly up-regulated (685) and down-regulated (209) genes for the three body segments (HvsLD50 & LHD55) (Fig. 2J,K). Upregulated genes belong to KEGG pathways: FoxO signaling, mTOR signaling pathway, Jak-STAT signaling pathway and glutathione metabolism^1,2,5^. Common downregulated DE genes affect KEGG pathways: metabolic pathways, FA biosynthesis, and amino acid (AA) biosynthesis. Similarly, there is an abundant number of tissue-specific DE genes associated with LH shift. A similar number of unique DE genes are upregulated and downregulated in heads (up=756; down=1138) and thorax (up=661; down=1018), while there is a lower number of DE genes in abdomen (up=311; down=685).

### Transcriptional analysis identifies key mediators of metabolic changes during shift

Organisms adapt to changes in energy by mobilizing stored nutrients and altering catabolic or anabolic processes to provide energy for survival^2^. At D50, before shift, changes in metabolism between H and L diets are mediated by increased expression of a limited number of common DE genes (HvsLD50): 1-Acylglycerol-3-phosphate O-acyltransferase 2 (Agpat2) and Diacylglycerol O-acyltransferase 2 (Dgat2), which have roles in triacylglycerol synthesis and droplet growth (Fig. 3A)^38^. HL shift results in using lipids as an energy source, as well as a decrease in lipid synthesis. This is reflected by reduced expression of common DE genes involved in lipid metabolism: Agpat2, Dgat2, yippee interacting protein 2 (yip2), Bubblegum (Bgm) Acetyl Coenzyme A synthase (AcCoAs), and Jabba (Fig. 3A, and not shown)^40^,^41^. while upregulation of these DE genes is linked to LH shift. The levels of Sterol regulatory element-binding protein (SREBP), which affects transcription of a variety of genes involved in de novo lipogenesis show an increasing trend during LH shift (Fig. 3A)^30,42^. LH shift is associated with reduced expression of Acetyl-CoA carboxylase (ACC), the rate-limiting substrate for FA synthesis (Fig. 3L). Inhibition of ACC or mitochondrial β-oxidation prevents dietary restriction (DR)-associated lifespan extension^33^. Lip3 catalyzes breakdown of TAG into free fatty acids (FFA) to be used as an energy source and its levels are strongly upregulated in abdomens and thoraces of HL shifted flies, and downregulated in LH flies^43^. Tissue-specific changes in metabolism are illustrated by different levels or trends in gene expression associated with shifts; for instance, Lip3 levels are increased in heads at D50, but are not affected by shifting, while in thorax the levels are lower in H than in L flies, and LH reduces its levels. In abdomen, HL is associated with an increase and LH with decrease in Lip3. Transcription of doppelganger von brummer (dob), a lipid droplet-associated TAG lipase, decreases in thorax and abdomens of HL shifted flies, while the levels are lower and constant in heads (Fig. 3L)^44^.

**Figure 3:**
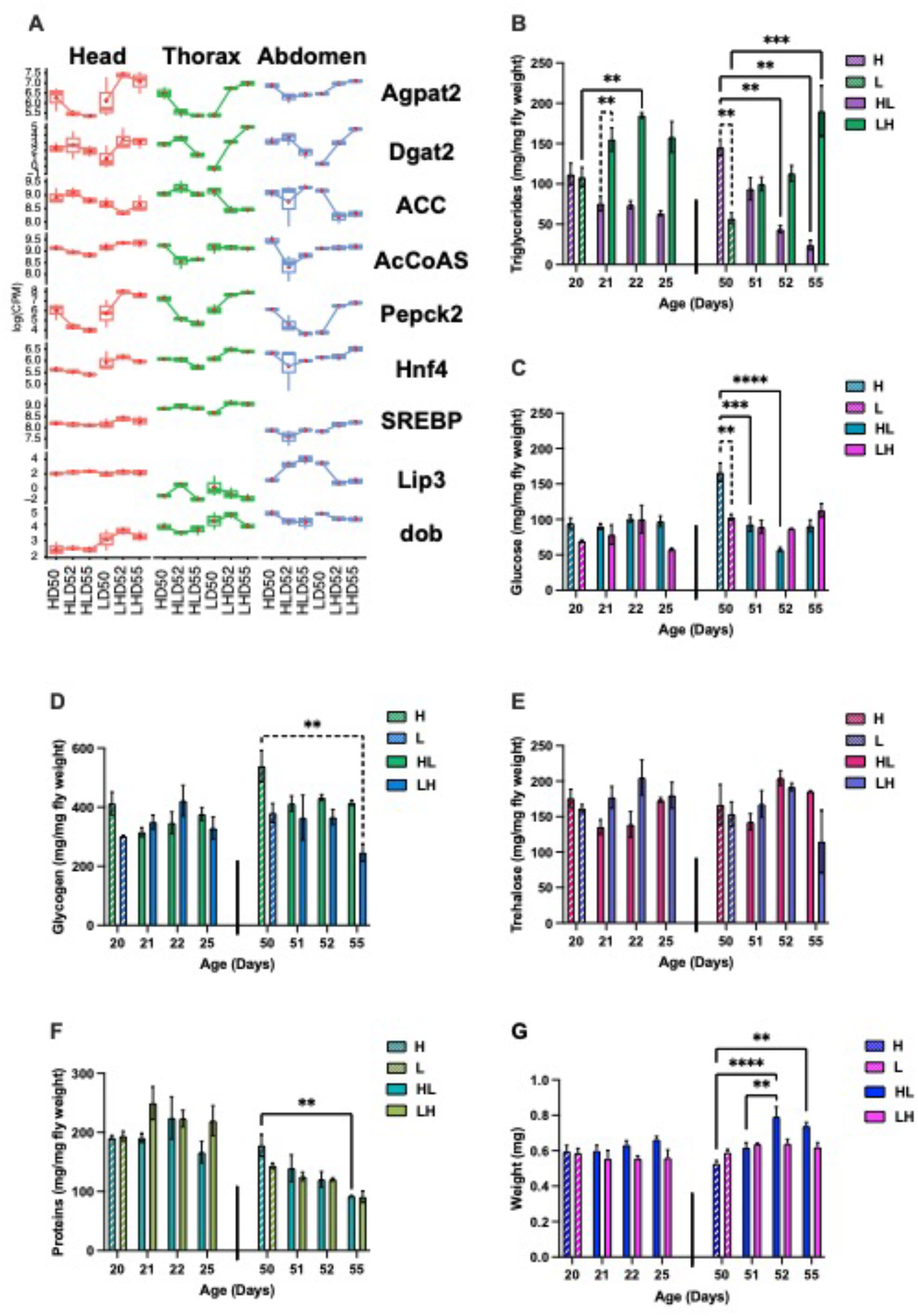
Age-dependent effects of shift on metabolism: **A)** Tissue-specific differentially expressed gene analysis of count data of Agpat2 (1-Acylglycerol-3-phosphate O-acyltransferase 2), Dgat2 (Diacylglycerol O-acyltransferase 2), ACC (Acetyl-CoA carboxylase), AcCoAS (Acetyl Coenzyme A synthase), Pepck2 (Phosphoenolpyruvate carboxykinase 2), Hnf4 (Hepatocyte nuclear factor 4), SREBP (Sterol regulatory element-binding protein), Lip3 (Lipase 3) and dob (doppelganger von brummer) during HL and LH shift. Levels of triglycerides **(B)**, glucose **(C)**, glycogen **(D)**, trehalose **(E)**, protein **(F)**, and weight **(G)** in male flies on day 20 and 50 before shifting, and 1, 2, and 5 days after shifting from H to L diet (HL) or L to H diet (LH). Error bars represent SEM. Data from 20 and 50 days were analyzed separately using two-way ANOVA. Post-hoc analysis was conducted using Tukey’s test, correcting for multiple comparisons. Results represent means ± SE of 3 biological replicates containing 10 flies per replicate. p: 0.033 (*), 0.002 (**), 0.0002 (***),<0.0001 (****).

### Flies adapt their metabolism to nutrient shifting at both young and old ages

To probe metabolic adaptation to H and L diets before and after the shifts on the organismal level, we determined the levels of triglycerides (TAG), glucose, glycogen, trehalose, and proteins in homogenates of whole male flies. These are important in metabolism as flies store excess energy as TAG in lipid droplets, and use carbohydrates as an energy source. Accordingly, these levels will alter depending on the nutritional status of the organism. To determine how age affects metabolic adaptations, we analyzed metabolism in 20- and 50-day-old flies aged on either a H or L diet (HD20; LD20; HD50; LD50). Subset of flies were then shifted to opposite diets, and metabolism was measured 1, 2, and 5 days after shifting (HLD21, HLD22, HLD25, LHD21, LHD22, LHD25) and the same for D50 (Fig.1A). The most dramatic changes are observed in TAG levels (Fig. 3B). At D20, flies on constant H or L diets have the same TAG levels indicating that young flies have metabolic resilience and easily adapt their TAG metabolism to different diets. However, by D50, aging on a constant H diet increases and aging on a constant L diet decreases TAG levels leading to an almost 2-fold difference between HD50 and LD50. This is mediated by increased expression of Agpat2 and Dgat2. At D20, HL shift leads to decreased TAG levels to about 60% of D20 5 days after shift. At D50, HL shift decreases TAG levels to about 20% of starting levels 5 days post-shift (HLD55) (Fig. 3B). LH shift increases the TAG levels at both ages, but is more striking at D50 as illustrated by a three-fold increase at D55. Our data show that 50D-old flies can immediately adapt their metabolism to nutrient conditions, consistent with immediate effects on mortality rate and HR (Fig. 1E,H,K,L).

Flies use glucose as a major source of energy, which can be stored as glycogen, or trehalose. At 20D, levels of glucose, glycogen, and trehalose are not different in flies on a L or H diet. At D50, levels of glucose are higher in HD50 flies, which is mediated by increased levels of Pepck2, the rate-limiting enzyme of gluconeogenesis^45^. HL shift decreases and LH increases Pepck2 levels affecting glucose levels within 24 hours (Fig. 3A,C). A similar trend in gene expression in all tissues was observed in Hepatocyte nuclear factor 4 (Hnf4), which mediates changes in glucose and lipid metabolism (Fig. 3A)^45^. HLD50 shift leads to a decreasing trend in glycogen levels (Fig. 3D), which may be due to glycogen’s role in supporting the high energy requirements of flight muscle. At D50, levels of trehalose are similar on H and L diets and shifts have no effect on trehalose levels (Fig. 3E). At D20 the levels of proteins are the same on L and H diets regardless of shift. At D50, HL shift decreases protein levels consistent with findings that CR decreases protein synthesis (Fig. 3F)^1,5^. The body weight of flies on H and L diets was similar at D20 (Fig. 3G). At D50, LH shift had no effect on weight, but HL increased weight.

### Tissue-specific gene trends mediate metabolic and physiological adaptation to diet shifts

To identify pathways underlying metabolic/physiological adaptation in each tissue, we analyzed tissue-specific gene trends using the degPattern function from the “DEGreport” package^31^. Changes in heads associated with diet shifting are mediated by ten unique gene trends, the thorax by nine, and the abdomen by six (Fig. 4A,C,D). Adaptive response to available energy is orchestrated by key hormones including DILPs, Akh, and steroid hormones, which integrate inter-organ signals leading to changes in metabolism and behavior^1^. Accordingly, in heads dramatic metabolic changes associated with HL shift are reflected by decreased levels of members of the IIS family and longevity pathway. (Fig. 4A,B). In heads, increased expression of several genes, including, Neuropeptide F receptor, Takeout, and Sodium/solute co-transporter-like 5A11 (SLC5A11) regulate behavioral changes associated with a sudden drop in energy^46^. In contrast, LH shift is associated with opposite gene trends and a decrease in those pathways. A substantial number of DE genes in head are also involved in ribosome and cytoplasmic translation. While expression of these genes changes slightly in HL shift, it increases dramatically in LH-shifted flies (Groups 6,8) (Fig. 4A,B). KEGG pathways include ribosome, endocytosis, proteasome, and apoptosis. The gene functional classification enrichment score for ribosomes is 16.21 and includes 47 ribosomal proteins. The brain, thorax, and abdomen adapt rapidly to decreases in available energy. HL shift increases and LH decreases DE genes in pathways associated with energy production, marked by enrichment of KEGG pathways such as oxidative phosphorylation in head (group 4) (p=1.8E-9), thorax (group 6) (p=1.6E-11), and abdomen (group 3) (p=2.6E-10).

**Figure 4:**
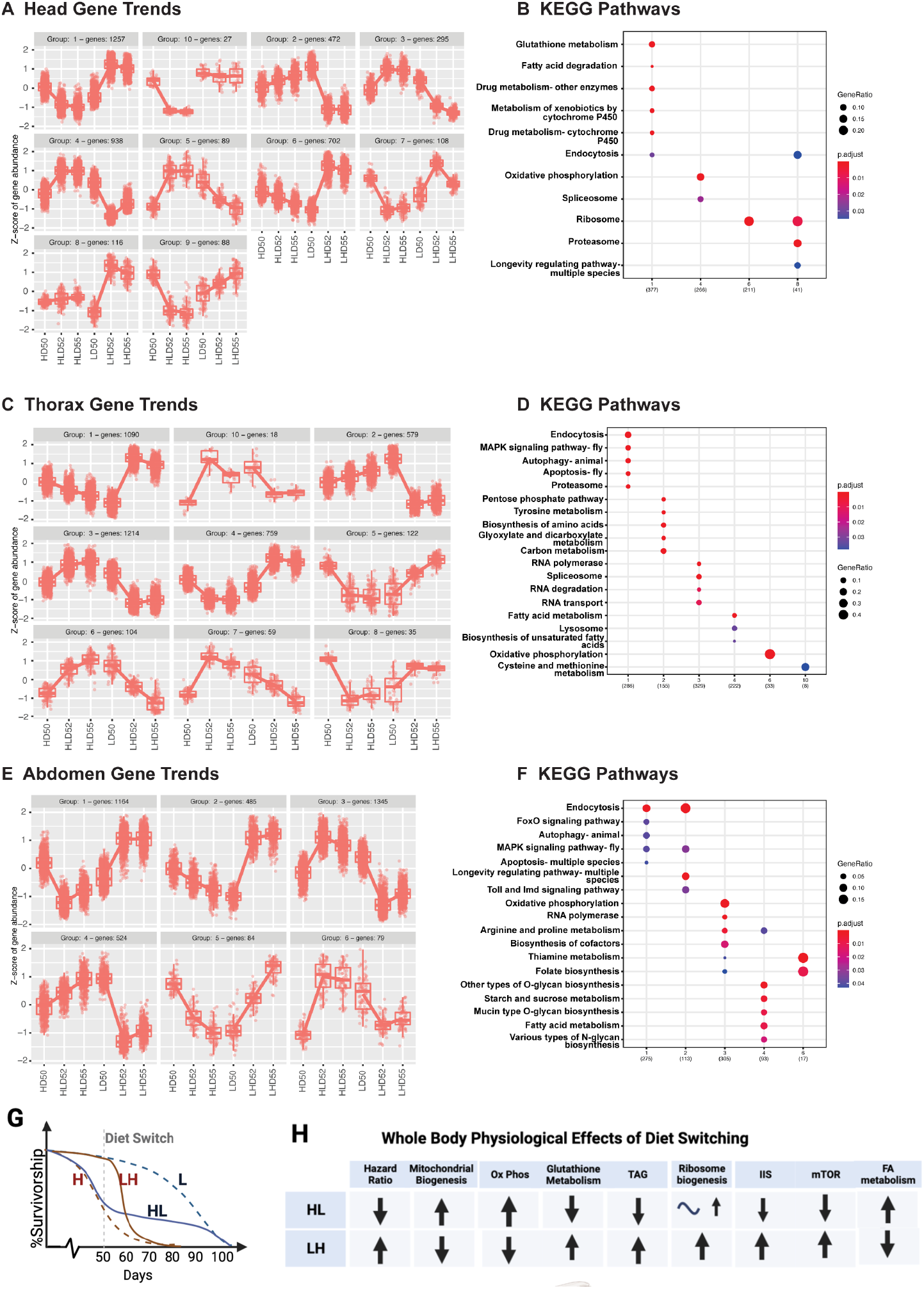
Tissue-specific and whole body physiological changes associated with diet switch at later age: Tissue-specific DE gene trends in Head **(A)** (ten groups), Thorax **(C)** (nine groups), and Abdomen **(E)** (six groups) associated with high (H) or low (L) diet on D50, or high to low (HL) or low to high (LH) shift on day two (D52) and five (D55) days after the HL (HLD52, HLD55) or LH (LHD52, LHD55) shift. **B**,**D**,**E)** KEGG functional enrichments pathways of Head (B), Thorax (D), and Abdomen (E) specific gene trends associated with diet shifting described in A, C, and E. **G, H**) Male flies rapidly adapt their whole body physiology to late life HL or LH shifts. Diet switch leads to longer lifespan in HL and shorter lifespan in LH shifted flies (G). H). Physiological adaptation associated with shifts leads to reduced (HL) or increased (LH) hazard ratio, glutathione metabolism and TAG levels, respectively. There is an increase in mitochondrial biogenesis, oxidative phosphorylation and fatty acid (FA) metabolism in HL shifted flies, while opposite is found in LH shift. In response to LH shift flies dramatically increase transcription and ribosome biogenesis, while a smaller increase is associated with HL shift.

While diet shifts clearly affect metabolism in all tissues, there is a difference in the trend of DE genes associated with HL or LH shift. HL shift leads to an increase in oxidative phosphorylation genes within 2 days and stays upregulated in heads and abdomens. In the thorax, there is a liner increase or decrease in oxidative phosphorylation genes that occurs after HL and LH shift, respectively (Fig. 4A,C,D). Additional changes during shifts are associated with mitochondrial function, and defense against ROS. In heads of HL shifted flies, upregulated DE genes belong the mitochondrion (p=6.7E-26), including mitochondrial translation (p=9.2E-16,), and ATP metabolic processes (p=2.7E-5), (group 4) (Fig. 4A). In contrast, LH shift is linked to decreased DE genes associated with mitochondrial function. HL shift decreases generation of ROS leading to a decrease in the glutathione metabolism pathway (p=3.2E-13), while LH shift increases generation of ROS leading to an increase in expression of genes in the glutathione metabolism pathway (GstT2, GstS1, GstE12) (Fig. 4 A,B). A similar decrease is observed in KEGG pathways indicated by an enrichment of pathways involved with metabolism of xenobiotics by Cytochrome P450.

The thorax is mostly built of muscles that have large energy requirements for flying; consequently, a HL shift leads to increases in mitochondrion (p=5.8E-9). The highest enrichment scores are in mitochondrial translation, mitochondrial small and large ribosomal units, ribosome biogenesis, and nucleocytoplasmic transport (Fig. 4C,D). A decreasing trend in HL and an increasing trend in LH shift (groups 1,4) belong to KEGG pathways: endocytosis, MAPK signaling pathway, apoptosis, proteasome, and FA metabolism. In contrast, LH shift decreases expression in genes (group 2) related to the pentose phosphate pathway and tyrosine metabolism (Fig. 4C).

Similarly, there is an enrichment of mitochondrion (p=5.3E-7) in abdomens of HL shifted flies. In abdomen, upregulated pathways support growth reflected by enrichment of folate biosynthesis, RNA polymerase, and ribosome biogenesis. The opposite is true of HL shifts, which decreases pathways related to growth like FoxO and MAPK signaling, as well as endocytosis and longevity-regulating pathway (groups 1,2,5) (Fig. 4E,F). Some upregulated genes stimulate proliferation of the germline stem cells including TGF-β signaling pathway (Smad), Notch, and Hippo signaling. Increased expression of Merlin (Mer) supports proliferation of germline and somatic cyst stem cells in *Drosophila* testis^47^, and is consistent with H diet increasing proliferation of male germline and intestinal stem cells in female flies^48^.

### KEGG Pathways across all samples

Functional enrichment analysis across all tissues and time points show significant changes in the components of the nutrient-sensing, stress response, and the signaling pathways that regulate metabolic homeostasis (Fig. 4G). KEGG pathways include longevity regulating pathway, endocytosis, ribosome, oxidative phosphorylation, metabolism and drug metabolism (Fig. 4A-D-G).

## Discussion

This study confers an immediate and long-term beneficial impact of shifting male flies from H to L diet late in life. Previous studies show that shifting flies at young or middle age from standard to CR diet results in immediate decrease in mortality rate and leads to lifespan extension, however, how late in life shift has beneficial effect remained unexplored^19,49^. Our data show that HL shift immediately lowers fly mortality rate and short-term hazard ratio (HR) below that observed in flies on a constant L diet, which within a few days becomes the same as flies on a constant L diet. We suggest that this short-term lower HR results from adaptation of male flies aged on a H diet for 50 days to cope with high nutrient levels. This adaptation is characterized by increased investment in protection from ROS production, as supported by increased expression of glutathione genes in HD50 flies. Flies on a high sucrose diet (HSD) have increased oxidative stress leading to increased expression of genes involved in ROS response^39^. While HSD differs from our H diet in that it has high sugar low protein content, it similarly upregulates genes involved in stress response. Thus, flies on a H diet are overprotected after HL shift, and also experience an immediate decrease in burden associated with a H diet. This is evidenced by the downregulation of the pathways related to redox status, oxidative stress, and detoxification within 5 days after shift. Studies in rodents, non-human primates, and humans indicate that CR results in metabolic adaptation associated with less oxidative damage to DNA^2,14^. Furthermore, increased TAG levels in male flies on H diet could be potentially beneficial by providing energy in flies shifted to L diet. This is consistent with the work showing that best survival in mice subjected to CR was observed in mice that preserved their fat mass longer under CR suggesting that preservation of fat could have a protective role^50,51^. ^52^

In contrast, we found that a LH shift leads to an immediate increase in HR, which is transiently higher than in male flies on a constant H diet. Flies aged on L diet adjust their physiology by downregulating growth and produce low levels of ROS. However, when shifted to a H diet, there is a sudden increase in energy acquisition, which is immediately used for grow and energy storage. These processes result in high production of ROS, which flies aged on a L diet have not adapted to. Due to lifelong, low levels of ROS, flies experiencing a sudden burst in metabolic activity may be at higher risk of death at the onset of a H diet, as was supported by our data. The number of DE genes associated with an LH shift is considerably higher compared to an HL shift. Particularly striking is an increase in the glutathione family, cytochrome p450, ribosome biogenesis, genes involved in growth, proliferation, and IIS. Upregulation of genes involved in ribosome biogenesis and Cytochrome p450 was found in female heads reared on HSD^39,53^.

Transferring fully fed young female and male to a diet with 35% less yeast and sugar showed that age-specific mortality depends only on current diet and there are no effects of previous diet^19^. A different group suggested that the increase in mortality observed in female flies after switch from CR to full-fed condition was a result of the cost associated with a nutrient-rich environment and hidden cost of DR^54^. However, another report showed that female flies re-fed after being dietary restricted have increased reproduction post-switch resulting in similar lifelong egg production as fully fed females without effect on survival, possibly^29^. Our experiments were designed to probe effects of late-life CR on male flies fed H diet and therefore differ from experiments where CR was applied to flies on a standard diet, or *ad libitum* (AD) fed rats or mice. However, out data suggest that in male flies sudden exposure to extreme abundance of energy, its utilization, release of ROS, all lead to dramatic transcriptomic activity that is costly for male flies.

Flies preserve metabolic homeostasis by adapting their metabolism to available nutrients^45^. The major metabolic change associated with HL shift is use of lipids instead of glucose as the major energy source. This is consistent with metabolic profiles of heads and thoraces of satiated and fasted flies, which show similar shift in energy utilization from glucose in satiated flies to ketone bodies generated from lipids in fasted flies^55^. Our transcriptomic data support both increased FA synthesis and degradation resulting in ketone production that is used as an energy source. Similarly, CR in flies leads to both lipolysis, due to use of lipids as energy source, as well as lipogenesis, which is required to replenish TAG used as energy source^33^. Mice exposed to CR have a brief period of increased FA synthesis in adipose tissue followed by a prolonged period of whole-body fat oxidation within a day. This metabolic adaptation balances increased FA oxidation and requirements to replenish lipid storage^56^ and is mediated by a several fold increase in transcription of key enzymes FA synthase and acetyl-CoA carboxylase several hours after food uptake, which drops below levels observed in AL mice before feeding. Such a metabolic shift reduces ROS production and contributes to benefits of CR. We observed an increase in biological processes associated with FA metabolism such as very long-chain FA biosynthetic processes, membrane lipid metabolic processes, and FA elongation in thoraces and abdomens of HL-shifted flies. Integration of the transcriptional response and metabolic changes is key to further understanding the mechanism of metabolic adaptation associated with diet shifting and require further metabolome analysis.

Overall, our work describes the effects of HL and LH diets shift on survivorship, mortality rate, metabolism, and tissue-specific whole-genome transcription of wild type *CS* male flies at old age. This information may also help us understand how middle age/older humans on high calorie diet could improve quality of life and longevity by altering factors such as overall food intake. As the incidence of metabolic syndrome increases with age, delineating age-dependent treatments and the potential of late life CR application, as well as mechanisms of CR are vital. Studies in flies, mice, and non-human primates show that CR can benefit organismal health without affecting survival^4,57,58^. Therefore, late-life diet shift in obese humans could have remarkable beneficial impact on health even if lifespan is not affected.

## Supporting information

Supplemental Information: Supplemental Figure Legends and Tables Titles.

## Acknowledgments

We thank Drs. Geneva Hargis, Gordon Carmichael and Kavitha Kannan for their helpful comments and suggestions. Figures 1A, 4G and 4F were prepared using BioRender. This work was supported by grants from the National Institute of Health: RO1AG059586, R01AG059586-03S1, the University of Connecticut (UConn) Claude D. Pepper Older Americans Independence Center (P30-AG067988) to B.R.; U24 HG009889, and R35 GM118140 to BRG; R01 NS106844 and R01 NS120556 to J.Y.H.L; T32HG010463 to B.J.H. Rogina is a recipient of a Glenn Award for Research in Biological Mechanisms of Aging.

## Author Contributions

Conceptualization, B.R.; Methodology and Investigation, M.L., J.M., K.M., D.M., D.M., S.O., B.R.G., J.Y.H.L., B.R.; Writing – Original Draft and Figure Creation, M.L., B.J.H., K.M., J.Y.H.L., B.R.; Writing – Review & Editing, Supervision, M.L., B.L.H., K.M., S.O., J.Y.H.L., B.R. Most authors contributed to the editing and proofreading of the final draft.

## Methods

### Fly strains, maintenance and diet

The standard wild-type *Canton-S (CS)* line was obtained from the Bloomington Stock Center (Stock number 1). *CS* flies were reared on food containing 25 mg/mL tetracycline for 3 generations to eliminate *Wolbachia*, followed by several generations in tetracycline-free food. Standard laboratory corn media was used to grow *CS* fly grandparents and parents from which cross was set to for F1 fly collections^22^. To avoid any effects of larval density on longevity, we had the same number of flies and time each group of flies stay in one vial starting with grandparents. Each cross for grandparents and parents: 10 virgin female and 9 male flies ages 4–10 days were crossed and passed to a new vial with corn media every 2 days. F1 progeny was collected every 24 hours. 25 male and 25 female flies were kept on a L or H food per experimental design. Flies were maintained in a humidified temperature-controlled environmental chamber at 25°C (Percival Scientific) on a 12-hour light:dark cycle with light on at 6 AM. Standard laboratory corn media as well as food marked as L = 0.5N and H = 3.0N were used^22^. The two food levels are standardized as 1.0X being the food that has 100 g/L of sucrose (MP Biomedicals, Inc), 100 g/L of brewer’s yeast (MP Biomedicals, Inc) and 20 g/L of agar^22,24^. Detailed food preparation and maintenance were described previously^22^.

### Lifespan Studies

Lifespan studies were performed using 10 to 12 groups (vials) of 25 male and 25 female flies placed in each vial, which were collected within 24 hours following eclosion as described above and maintained in plastic vials containing high or low-calorie diet, as described below. Flies were kept in a humidified, temperature-controlled incubator with 12/12-h on/off light cycle at 25 °C. While males were aged together with female flies, here we present data for only male flies to avoid confounding effects of female reproduction, which are strongly affected by nutrients and need to be further examined carefully. The number of flies in each survivorship study is listed in Supplemental Tables 1,2,6,7. Two different shifting experiments were performed. Experiment 1: Two groups of *CS* flies on a high-calorie (H) or low-diet (L) for their whole lifespan, and three groups on L and three groups on H shifted to opposite food at ages 20, 50, or 60 days. Flies were passed every day from day 1 and the number of dead flies were counted. Experiment 2: The experimental groups: two groups, *CS* flies living their whole life on a H or L diets, and five groups that began on H from birth and were moved to L at either 10, 20, 30, 40 or 50 days. Five additional groups of flies were kept on L diet from birth and shifted from L to H at either 10, 20, 30, 40 or 50 days. They were passed every 2 days up to age 10 days and every day after that and the number of dead flies were counted.

### Biochemistry

*Canton-S (CS)* flies were collected and aged as described above on L or H diets. At ages 20 and 50, subgroups of flies were transferred to the opposite diet. Flies kept their whole life on L or H were used for analysis at age 20 or 50 days. Groups of flies transferred to opposite diet at 20 days were used for analysis at ages 21, 22, and 25 days. Similarly, flies that were transferred at 50 days to the opposite diet were used for analysis at ages 51, 52, and 55 days. Flies were sorted on CO^2^ by sex between 9:00AM – 9:30AM. 3 biological replicates of 10 male flies, were anesthetized on CO^2^, weighed, homogenized in cold homogenization buffer (0.01M KH_2_PO_4_, 1 mM EDTA pH 7.4). The tube was spun down at 2000 rpm for 2 minutes at 4°C. 25 μl of each homogenate was aliquoted into three wells of each of six 96-well plates. The plates were kept on dry ice allowing homogenates to be frozen immediately. The plates were covered and kept at -80°C until quantification of glucose, glycogen, trehalose, triglyceride, and protein. Before quantification, the plates were taken from -80°C and left to warm up to room temperature for 15 minutes. For glucose Glucose Assay Kit (Sigma GAG020, PGO Sigma P7119) was used as described previously^59^. For glycogen, the procedure was the same as glucose except amyloglucosidase (Amyloglucosidase from *Aspergillus niger*, Sigma 10115) was added to each well in addition to the other enzyme. For trehalose, the procedure was the same as glucose except the samples were incubated with trehalase before adding PGO (Trehalase from porcine kidney, Sigma T8778). Protein was determined using Total Protein Kit, Micro Lowry,Peterson’s Modification: (Sigma TP0300). The triglycerides were determined using Serum Triglyceride Determination Kit (Sigma TR0100; Glycerol Standard Solution Sigma G7793)^59^.

### Hazard Ratio (HR)

Data binning: Survival data were preprocessed using custom python scripts and binned in groups of 10 days. The hazard ratios were calculated on binned data from flies, which were shifted from one diet to another (HL or LH) at 20, 50, and 60 days and compared to flies which were kept on constant H or L diets. Custom scripts in R (Version: 4.1.2) were written to calculate hazard ratios using the cox proportional hazard regression model. The model was made, and hazard ratios calculated using the survival library (Version: 3.2-13). To generate a summary table of the cox model tabcoxph function from the tab library (Version: 5.1.1) was used. To check for the proportional hazard assumption for a cox regression model scaled Schoenfeld residual test was performed and visualized for the covariates using the R libraries survival and survminer (Version:0.4.9) respectively.

|  |  |
| --- | --- |
| Variable | variable name |
| Beta (SE) | Coefficient (standard error) |
| HR (95% CI) | Hazard Ratio (confidence Intervals) |
| P | P values |

HR is calculated from Beta by taking its exponent value. Values of Beta are determined by the cox regression model. Log-rank test was used to calculate the P values.

### RNA Extraction

For RNA-Seq experiments flies were collected within 24 hours following eclosion. 25 male and 25 female flies were placed in each vial and maintained in plastic vials containing high (H) or low (L) calorie diet, and transferred to opposite diet at age 50 days. At ages 50 (before shift), and 52 and 55 (2 or 5 days after shift) days, the flies were separated by sex on CO_2_ and immediately frozen at the same time of the day. Frozen male flies were dissected into heads, thoraces, and abdomens on a glass plate placed on top of a frozen block to keep tissues frozen. Total RNA was isolated from the heads, thoraces, and abdomens of 3 biological replicates with more than 50 flies in each replicate using Trizol as described^60^.

### Library preparation and sequencing

One μg of each RNA sample was used to prepare RNA-seq libraries with the Illumina Tru-Seq Stranded mRNA Library Preparation kit (cat # 20020594) with the IDT for Illumina TruSeq RNA UD Indexes set (cat # 20040871). The libraries were pooled and sequenced on one lane of a NovaSeq 6000 S4 flow cell generating 100bp paired-end reads yielding an average of 80 million reads per sample and a minimum of 20 million reads per replicate.

#### mRNA-Seq analysis

The quality of the sequencing data was evaluated with FastQC (v0.11.9) and MultiQC (1.9). Sequence reads were mapped to the Drosophila genome (GSE97233) with STAR (version) (2.71a) and the resulting BAM files were used to generate gene counts with featureCount using the uniq-counting mode. Analyses of sample relationship and differential expression were performed with DESeq2^32^. One sample, head LD50 (TLD50) biological sample (#39), had a notably high percentage of unmapped and unassigned reads and was therefore excluded from the downstream analysis. To identify specific pathways associated with diet shift, we performed functional enrichment analysis using the DAVID database. Pathway analysis and identification of networks associated with shifting flies were also carried out by Gene Ontology (GO) and the KEGG.

### Statistical analysis

Data from 20- and 50-day-old flies were analyzed separately using two-way ANOVA. Post-hoc analysis was conducted using Tukey’s test, correcting for multiple comparisons using GraphPad Prism 9.4.1. Results represent means ± SE of 3 biological replicates containing 30 flies per replicate. p: 0.033 (*),0.002 (**), 0.0002 (***), <0.0001 (****). Longevity data were censored for early mortality (1 - 10 Days) and analyzed by log-rank tests using the JMP16. program.

