## Supplemental Information: Supplemental Figure Legends and Tables Titles. for "Late- life shift in caloric intake affects fly longevity and metabolism"

**Supplemental Information**
**Supplemental Figure titles and legends**

**Supplemental Figure 1: Shifting diets has immediate effects on male lifespan and mortality**: Survivorships (A,B) and mortality rates (C-L) of male flies shifted from H to L at day 10 (HLD10), day 20 (HLD20), day 30 (HLD30), day 40 (HLD40) or day 50 (HDL50) (C-G) or from L to H diet on day 10 (LHD10), day 20 (LHD20), day 30 (LHD30), day 40 (LHD40) or day 50 (LHD50) (H-L). Survivorships curves and mortality rate were analyzed by long-rank test JMP16 program.

**Supplemental Figure 2: Shifting diet induces striking tissue-specific changes in transcriptome: A)** A sample-to-sample distance heatmap shows the driving force for distance in each tissue type or specific tissues, marked Abdomen, Thorax, or Head, of differentially expressed (DE) genes at high (H) or low (L) diets at day 50 (D50) before and 2 and 5 days after H to L shift. **B)** Heatmap of all DE genes in abdomen (A), head (H) and thorax (T) between flies on High or Low diets at day 50 (D50) before, and day 52 (D52), and 55 (D55), 2 and 5 days after shift.

**Supplemental Figure 3: Tissue-specific DE gene trends reveal tissue-specific biological processes:** Functional enrichment analysis of DE genes in head (A), thorax (B) and abdomen (C) associated with high (H) or low (L) diet on day 50, or high to low or low to high shift on day two and five days after shift.

**Supplemental Tables**
**Supplemental Table 1:** Effects of shifting *Canton-S* male flies from a high (H) to a low (L) calorie diet at 20, 50 and 60 days on longevity in comparison to longevity of flies on lifelong high calorie diet.

**Supplemental Table 2:** Effects of shifting *Canton-S* male flies from a low (L) to a high (H) calorie diet at 20, 50 and 60 days on longevity compared to longevity of flies on lifelong low calorie diet.

**Supplemental Table 3:** Hazard ratio (HR) of *Canton-S* male flies shifted at 50 (A), 60 (B) or 20 (C) days from a high (H) to a low (L) calorie diet (HL) or vice-versa (LH) and compared to hazard ratio of flies kept on lifelong high (H) or low (L) calorie diet

**Supplemental Table 4:** Effects of shifting *Canton-S* male flies from a high (H) to a low (L) calorie diet at 10, 20, 30, 40 and 50 days on longevity compared to longevity of flies on lifelong high calorie diet.

**Supplemental Table 5:** Effects of shifting *Canton-S* male flies from a low (L) to a high (H) calorie diet 10, 20, 30, 40 and 50 days on longevity compared to longevity of flies on lifelong low calorie diet.
