## Supplementary figures and images for "Late- life shift in caloric intake affects fly longevity and metabolism"

### Supplemental Figure 1

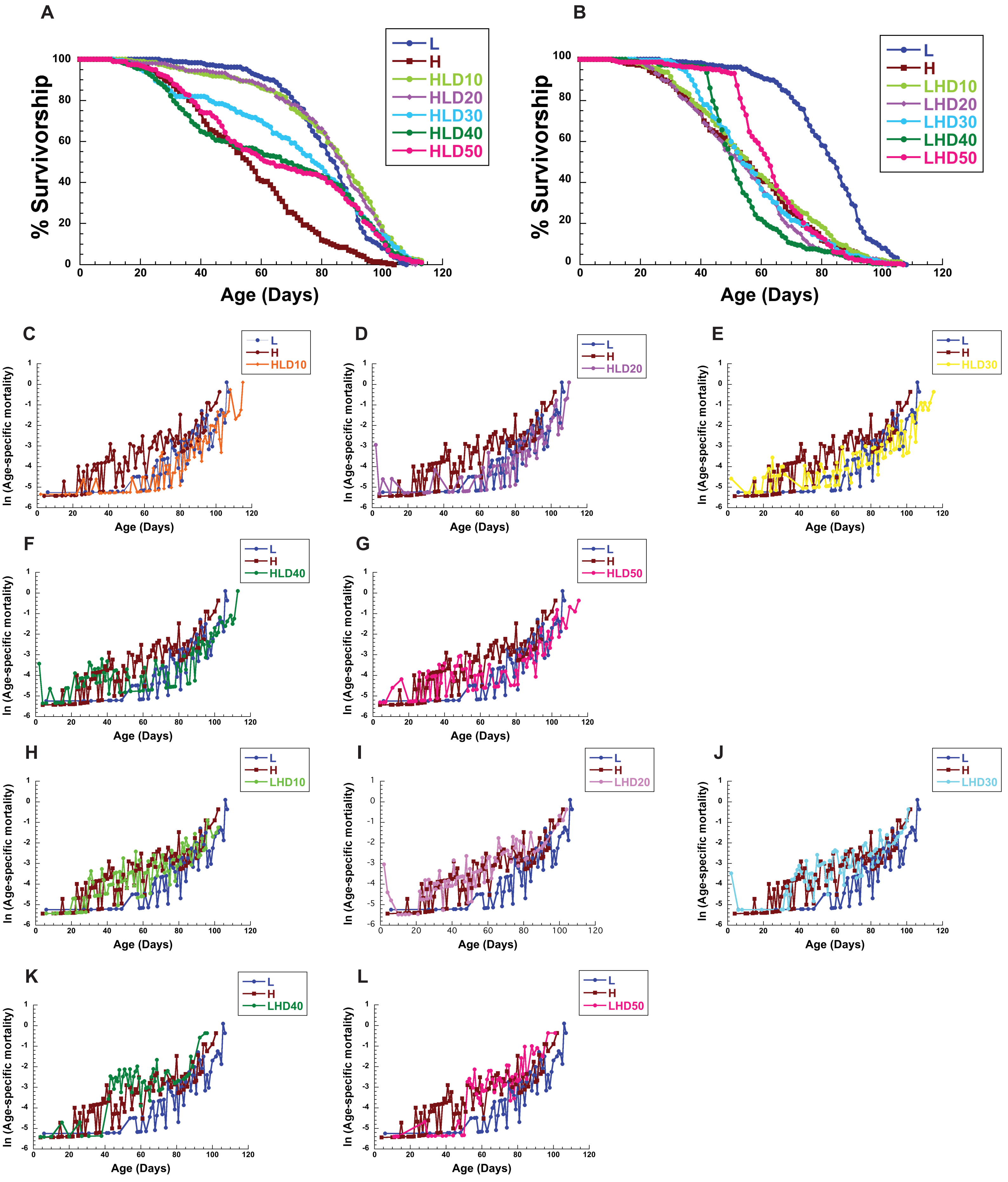

### Supplemental Figure 2

**A**

#### Sample Distances

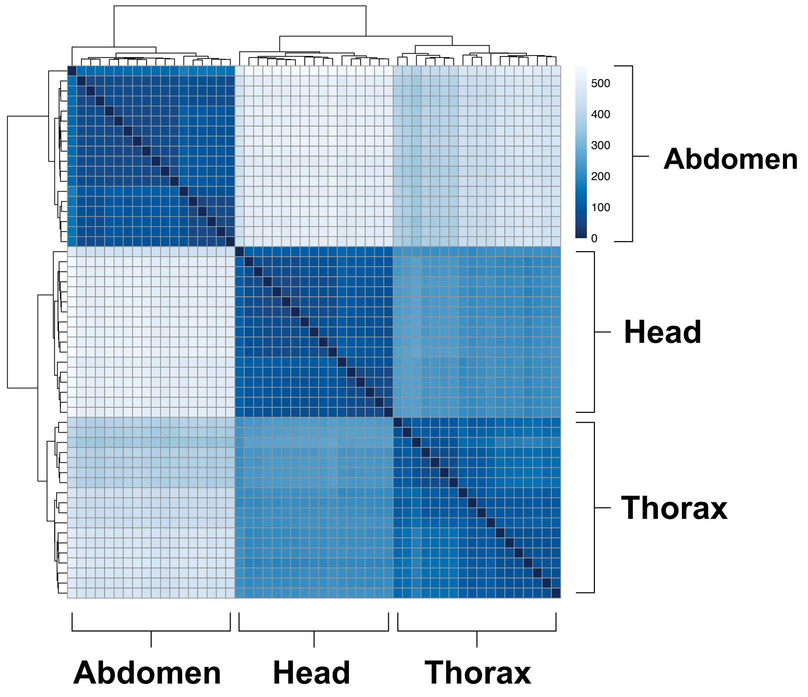


**B**

#### Heatmap of All Tissues

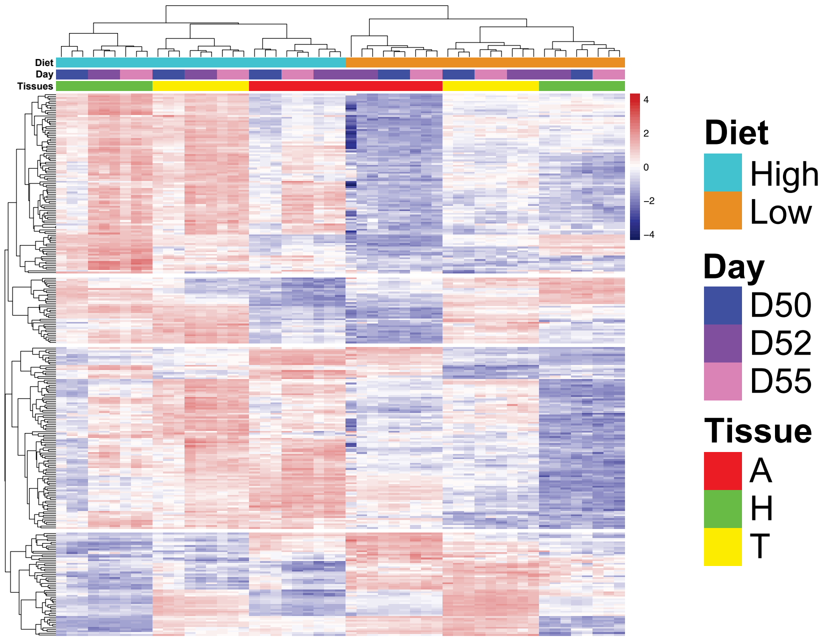
