## Supplemental Table 1 for "Late- life shift in caloric intake affects fly longevity and metabolism"

**Supplemental Table 1: Effects of shifting *Canton-S* male flies from a high (H) to a low (L) calorie diet at 20, 50 and 60 days on longevity compared to longevity of flies on lifelong high calorie diet.**

| **Food** | **Time on L Diet** | **N** | **Mean LS (% change)** | $X^2$ | **p** | **Maximal LS (% change)** |
|---|---|---|---|---|---|---|
| H | Lifelong | 385 | 41.2 | 703.0448 | <0.0001* | 72.8 |
| L | Lifelong | 411 | 86.7 | | | 107.2 |
| H | Lifelong | 385 | 41.2 (-93.4) | 383.7028 | <0.0001* | 72.8 (-39.7) |
| HLD20 | D20 | 226 | 79.7 | | | 101.7 |
| H | Lifelong | 385 | 41.2 (-17.7) | 21.1998 | <0.0001* | 72.8 (-39.7) |
| HLD50 | D50 | 222 | 48.5 | | | 101.7 |
| H | Lifelong | 385 | 41.2 (-4.6) | 3.1870 | 0.0742 | 72.8 (-26.6) |
| HLD60 | D60 | 216 | 43.1 | | | 92.2 (35.2) |

L = Low calorie diet
H = High calorie diet
HLD20 = flies shifted from a high to low calorie diet at day 20
HLD50 = flies shifted from a high to low calorie diet at day 50
HLD60 = flies shifted from a high to low calorie diet at day 60
*Statistically significant
