## Supplemental Table 2 for "Late- life shift in caloric intake affects fly longevity and metabolism"

**Supplemental Table 2: Effects of shifting *Canton-S* male flies from a low (L) to a high (H) calorie diet at 20, 50 and 60 days on longevity compared to longevity of flies on lifelong low calorie diet.**

| Food | Time on L Diet | N | Mean LS (% change) | $X^2$ | p | Maximal LS (% change) |
|---|---|---|---|---|---|---|
| L | Lifelong | 411 | 86.7 (52) | 703.0448 | <0.0001* | 107.2 (32) |
| H | Lifelong | 385 | 41.2 | | | 72.8 |
| L | Lifelong | 411 | 86.7 (57 | 636.7169 | <0.0001* | 107.2 (41) |
| LHD20 | D20 | 210 | 37.6 | | | 63.5 |
| L | Lifelong | 411 | 86.7 39) | 516.7181 | <0.0001* | 107.2 (42) |
| LHD50 | D50 | 231 | 52.6 | | | 61.7 |
| L | Lifelong | 411 | 86.7 (30) | 404.2337 | <0.0001* | 107.2 (36) |
| LHD60 | D60 | 215 | 60.3 | | | 68.2 |

L=Low calorie diet
H=High calorie diet
LHD20= flies shifted from a low to high calorie diet at day 20
LHD50= flies shifted from a low to high calorie diet at day 50
LHD60= flies shifted from a low to high calorie diet at day 60
*Statistically significant
