## Supplemental Table 3 for "Late- life shift in caloric intake affects fly longevity and metabolism"

**A) Hazard ratio (HR) of *Canton-S* male flies shifted at 50 days from a high (H) to a low (L) calorie diet (HL) or vice-versa (LH) and compared to hazard ratio of flies kept on lifelong high (H) or low (L) calorie diet.**

| Period (days) | HR H&HLD 50 | p | HR H&LHD50 | p | HR L&HLD50 | p | HR L&LHD5 0 | p |
|---|---|---|---|---|---|---|---|---|
| 50-60 | 0.99 | 0.96 | 1.53 | 0.005* | 1.75 | 0.09 | 2.87 | < 0.001* |
| 60-70 | 0.29 | 0.02* | 0.61 | 0.2 | 0.42 | 0.12 | 0.87 | 0.73 |
| 70-80 | 0.20 | 0.008* | 0.15 | 0.01* | 0.21 | 0.01* | 0.19 | 0.03* |
| 80-90 | | | | | 0.25 | 0.02* | | |

*Statistically significant

**B) Hazard ratio (HR) of *Canton-S* male flies shifted at 60 days from a high (H) to a low (L) calorie diet (HL) or vice-versa (LH) and compared to hazard ration of flies kept on a constant high (H) or low (L) calorie diet.**

| Period (days) | HR H&HLD60 | p | HR H&LHD60 | p | HR L&HLD60 | p | HR L&LHD60 | p |
|---|---|---|---|---|---|---|---|---|
| 60-70 | 0.66 | 0.28 | 2.02 | 0.002* | 0.8 | 0.56 | 2.67 | < 0.001* |
| 70-80 | 0.07 | 0.01* | 0.23 | 0.02* | 0.08 | 0.02 | 0.25 | 0.03* |

*Statistically significant

**C) Hazard ratio (HR) of *Canton-S* male flies shifted at 20 days from a high (H) to a low (L) calorie diet (HL) or vice-versa (LH) and compared to hazard ratio of flies kept on a constant high (H) or low (L) calorie diet.**

| Period(days) | HR H&HLD20 | p | HR H&LHD20 | p | HR L&HLD20 | p | HR L&LHD20 | p |
|---|---|---|---|---|---|---|---|---|
| 20-30 | 0.47 | 0.15 | 1.19 | 0.27 | 1.17 | 0.71 | 1.26 | 0.49 |
| 30-40 | 0.96 | 0.9 | 1.12 | 0.49 | 0.35 | 0.1 | 1.57 | 0.16 |
| 40-50 | 0.51 | 0.11 | 1.36 | 0.09 | 0.85 | 0.68 | 1.66 | 0.09 |
| 50-60 | 0.44 | 0.07 | 0.84 | 0.48 | 0.84 | 0.69 | 1.76 | 0.1 |
| 60-70 | 0.54 | 0.07 | 0.5 | 0.2 | 0.76 | 0.47 | 0.96 | 0.93 |
| 70-80 | 0.42 | 0.02* | 0.17 | 0.02* | 0.56 | 0.06 | 0.35 | 0.06 |

*Statistically significant
