## Supplemental Table 4 for "Late- life shift in caloric intake affects fly longevity and metabolism"

**Supplemental Table 4: Effects of shifting *Canton-S* male flies from a high (H) to a low (L) calorie diet 10, 20, 30, 40 and 50 days on longevity compared to longevity of flies on lifelong high calorie diet.**

| **Food** | **Time on L Diet** | **N** | **Mean LS (% change)** | $X^2$ | **p** | **Maximal LS (% change)** |
|---|---|---|---|---|---|---|
| H | Lifelong | 227 | 55.9 (-48.2) | 173.88415 | <0.0001* | 93.0 (-15.9) |
| HLD10 | D10 | 210 | 82.8 | | | 107.9 |
| H | Lifelong | 227 | 55.9 (-47.4) | 151.6646 | <0.0001 | 93.0 (-12.6) |
| HLD20 | D20 | 199 | 82.5 | | | 104.8 |
| H | Lifelong | 227 | 55.9 (-27.9) | 67.7344 | <0.0001 | 93.0 (-16.5) |
| HLD30 | D30 | 194 | 71.5 | | | 108.5 |
| H | Lifelong | 227 | 55.9 (-16.1) | 37.3879 | <0.0001 | 93.0 (-14.3) |
| HLD40 | D40 | 208 | 64.9 | | | 106.4 |
| H | Lifelong | 227 | 55.9 (-18.4) | 37.4033 | <0.0001 | 93.0 (-12.6) |
| HLD50 | D50 | 200 | 66.2 | | | 104.8 |

L=Low calorie diet
H=High calorie diet
HLD10= flies shifted from a high to low calorie diet at day 10
HLD20= flies shifted from a high to low calorie diet at day 20
HLD30= flies shifted from a high to low calorie diet at day 30
HLD40= flies shifted from a high to low calorie diet at day 40
HLD50= flies shifted from a high to low calorie diet at day 50
*Statistically significant
