## Supplemental Table 5 for "Late- life shift in caloric intake affects fly longevity and metabolism"

**Supplemental Table 5: Effects of shifting *Canton-S* male flies from a low (L) to a high (H) calorie diet at 10, 20, 30, 40 and 50 days on longevity compared to longevity of flies on lifelong low calorie diet.**

| Food | Time on L Diet | N | Mean LS (% change) | $X^2$ | p | Maximal LS (% change) |
|---|---|---|---|---|---|---|
| L | Lifelong | 189 | 81.8 (32.0) | 122.5170 | <0.0001* | 103.7 (10.2) |
| H | Lifelong | 227 | 55.9 | | | 93.0 (-10.9) |
| L | Lifelong | 189 | 81.8 (29.2) | 89.0624 | <0.0001* | 103.7 (8.6) |
| LHD10 | D10 | 226 | 57.9 (-41.3) | | | 94.3 (-9.5) |
| L | Lifelong | 189 | 81.8 (34.6) | 166.4088 | <0.0001* | 103.7 (15.8) |
| LHD20 | D20 | 240 | 53.5 (-52.9) | | | 86.9 (-18.7) |
| L | Lifelong | 189 | 81.8 (12.5) | 132.5256 | <0.0001* | 103.7 (12.5) |
| LHD30 | D30 | 194 | 57.5 (-42.3) | | | 90.4 (-14.2) |
| L | Lifelong | 189 | 81.8 (34) | 205.7889 | <0.0001* | 103.7 (17) |
| LHD40 | D40 | 222 | 54.1 (-51.4) | | | 85.2 (-21.2) |
| L | Lifelong | 189 | 81.8 (22.2) | 135.4157 | <0.0001* | 103.7 (14.9) |
| LHD50 | D50 | 221 | 63.6 (-28.6) | | | 87.9 (-17.5) |

L = Low calorie diet
H = High calorie diet
LHD10 = flies shifted from a low to a high calorie diet at day 10
LHD20 = flies shifted from a low to a high calorie diet at day 20
LHD30 = flies shifted from a low to a high calorie diet at day 30
LHD40 = flies shifted from a low to a high calorie diet at day 40
LHD50 = flies shifted from a low to a high calorie diet at day 50
*Statistically significant
